# Investigating T cell Recruitment in Atherosclerosis using a novel Human 3D Tissue-Culture Model reveals the role of CXCL12 in intraplaque neovessels

**DOI:** 10.1101/2024.02.14.580316

**Authors:** Laura Parma, Nadja Sachs, Zhaolong Li, Kevin Merchant, Nikola Sobczak, Bram Slütter, Lars Maegdefessel, Christian Weber, Johan Duchene, Remco T.A. Megens

**Affiliations:** Institute for Cardiovascular Prevention (IPEK), Ludwig-Maximilians University, Pettenkoferstrasse 8a & 9, 80336 Munich, Germany; German Center for Cardiovascular Research (DZHK), partner site Munich Heart Alliance, Pettenkoferstrasse 8a & 9, 80336 Munich, Germany; Department for Vascular and Endovascular Surgery, Klinikum rechts der Isar, Technical University Munich, Munich, Germany; Division of Biotherapeutics, Leiden Academic Centre for Drug Research, Leiden University, Einsteinweg 55, Leiden, the Netherlands; Department of Medicine, Center for Molecular Medicine, Karolinska Institute, Stockholm, Sweden; Cardiovascular Research Institute Maastricht, University of Maastricht, P. Debeyelaan, 6229HX Maastricht, the Netherlands; Munich Cluster for Systems Neurology (SyNergy), Munich, Germany

## Abstract

**Background:** Development of effective treatments for atherosclerosis requires new models that better predict the human immune response. Although T cells are abundant in human atherosclerotic lesions and play a key role in the pathogenesis, the mechanism involved in plaque infiltration remains ill defined.

**Methods:** We developed a three-dimensional tissue-culture model to study leukocyte recruitment to human atherosclerotic plaques. In this study, human atherosclerotic plaques obtained during carotid endarterectomy surgery were co-cultured with patient-matching T cells. Exogenous T cells were stained using a multi-factor staining strategy, which involved intracellular fluorescent cell tracker dyes combined with nuclear labels. Flow cytometry was used to assess the presence of the labeled cells within the plaques, and microscopic analysis was performed to examine their localization.

**Results:** Flow cytometry and microscopy cell-tracking analysis demonstrated that exogenous T cells successfully migrated into atherosclerotic plaques. Furthermore, infiltrated CD8^+^ T cells displayed a significant increase of CD69 expression, indicating their activation within the tissue. Blocking chemokine receptors, particularly CXCR4, significantly impaired T cell infiltration, demonstrating that exogenous CD8^+^ T cells invade plaques through chemotactic migration. Surprisingly, 3D microscopy combined with optical tissue clearing strategy revealed that CXCL12, the sole ligand of CXCR4, mainly accumulated in intraplaque neovessels. Single-cell RNA sequencing (scRNAseq) analysis further confirmed that endothelial cells from intraplaque neovessels were the primary source for CXCL12. Additionally, exogenous T cells were found within and in proximity to these neovessels, suggesting that the CXCL12/CXCR4 axis regulates T cell recruitment through intraplaque neovessels.

**Conclusions:** Overall, these findings shed new light on the mechanism of action of CXCL12 in atherosclerosis and demonstrated the potential of the model to advance our understanding of leukocyte accumulation in human atherosclerosis and assist in testing novel pharmacological therapies.

## 1. Introduction

Cardiovascular disease (CVD) is the number one cause of death worldwide, taking an estimated 18 million lives each year^1^. The most common underlying cause of life-threatening cardiovascular event is atherosclerosis, which culminates in plaque rupture, the most critical disease stage^2^. Classically, atherosclerosis has been viewed as a disorder of lipid deposition within the vessel wall of arteries and has therefore been treated focusing on lowering LDL cholesterol levels^3^. However, not all patients benefit from lipid-lowering therapies and therefore it is imperative to seek alternative treatments.

Contrary to what was traditionally suggested, that macrophages and foam-cells dominated the atherosclerotic landscape, recent results from single-cell RNA sequencing (scRNA-seq) studies, in which mapping of the whole inflammatory cell content of human atherosclerotic plaques has been carried out^4–6^, have revealed that atherosclerotic plaques are enriched for activated T cells. The latter suggests that T cells are critical drivers and modifiers in the pathogenesis of atherosclerosis^4, 7^. Moreover, recent studies established that atherosclerosis is characterized by an autoimmune component, which in humans is driven by autoreactive effector T cells^6, 8^. To precisely link T cell phenotypes to clinical outcomes and to generate novel treatment options, it is pivotal to unravel which stimuli drive T cells recruitment, activation, and interaction with other cell types in human atherosclerotic plaques.

Despite the considerable research effort conducted to understand the mechanisms driving the development and advancement of atherosclerosis in mouse models lacking apolipoprotein E (ApoE^-/-^) or low-density lipoprotein receptor (Ldlr^-/-^), translating the insights gained from murine T-cell immunology to the human context has proven to be a significant challenge. In fact, as a recent global call to action for cardiovascular disease (CVD) drug solutions highlighted, several interventions that demonstrated efficacy in preclinical animal models have ultimately proven to be insufficient predictors of effective interventions in human trials^9^.

Comparing the results from single cell studies in mouse atherosclerosis with recent studies exploring single cell phenotypes of T cells in human atherosclerosis, it became clear that there are substantial differences between T cells in mice when compared to humans^4, 5, 10–12^. For instance, only few groups of T cells were present in mouse aortas^10, 11, 13, 14^, while a significantly greater diversity of T-cell populations was found in human atherosclerotic plaques^4, 5, 15^.

To overcome these translational issues and to study the mechanisms involved in atherosclerotic plaque leukocyte recruitment, we developed a 3D tissue-culture model of human atherosclerosis using whole mount atherosclerotic plaques obtained from patients affected by carotid artery disease (CAD) during carotid endarterectomy (CEA) and T cells isolated from patient-matching blood samples. Our study demonstrates that isolated T cells (exogenous T cells) can be co-cultured with carotid plaque biopsies and their infiltration in the plaque can be studied quantitatively, using fluorescence-activated cell sorting (FACS) while their location can be mapped by combining a novel optical tissue clearing protocol with advanced microscopic imaging modalities. Application of the novel 3D tissue-culture model of atherosclerosis revealed that chemokines and their receptors play an essential role in T cell plaque recruitment and that blockade of CXCR4 on exogenous CD8^+^ T cells impairs their infiltration. Moreover, by using optical tissue clearing combined with multidimensional microscopy we identified the presence of CXCL12 (CXCR4 ligand) on the endothelial cells (ECs) lining intraplaque neovessels, as well as exogenous CD8^+^ T cells within and in their close vicinity, thereby proposing a new explanation for the detrimental role played by CXCL12 in atherosclerosis.

## 2. Material and Methods

### 2.1. Human Carotid Endarterectomies

Atherosclerotic samples were harvested during carotid endarterectomy surgeries at the Department of Vascular and Endovascular Surgery (Klinikum rechts der Isar, TUM) from patients with carotid artery disease (CAD) upon patients’ informed consent, as described previously^16^. Biopsies contained both the atherosclerotic plaque (advanced stage) and the adjacent region (early stage). The Munich Vascular biobank is approved by the local Hospital Ethics Committee (2799/10, Ethikkommission der Fakultät für Medizin der Technischen Universität München, Munich, Germany) and performs in accordance with the Declaration of Helsinki. Detailed information of the patients are reported in Supplemental Table 1. Plaques were processed differently for either single cell RNA sequencing or plaque culture, as described below.

### 2.2. Single cell RNA sequencing (scRNA-seq)

Plaque tissue preparation for single cell RNA sequencing was performed as previously described^17^.

### 2.3. Human plaque culture

For the plaque culture protocol, only the atherosclerotic plaque part of the biopsies were used. Plaque samples were dissected and divided into two parts of 1mm thickness. The plaques were placed on a wetted gelatin sponge raft (Spongostan standard, MS002, Ethicon) at the medium–air interface as previously described^18^ inside a Petri-dish (CC7682-3340, CytoOne). Advanced RPMI 1640 cell culture medium (12633-012, Gibco Life Technologies) supplemented with antimycotic antibiotic (15240-062, Gibco Life Technologies) and Penicillin-Streptomycin antibiotic (15140122, Gibco Life Technologies), containing freshly isolated patient-matching T cells was added to the culture and kept for 24 hours at 37 °C in a humidified 5% CO2 environment. After 24 hours of culture, plaque, sponges and culture medium were collected for further processing. One plaque slice, together with the sponge and the culture medium were analysed by means of flow cytometry. The second plaque slice was preserved in 2% PFA (8.18715.1000, Merck Millipore) and used for optical tissue clearing.

### 2.4. T cells isolation, labelling and treatment

Blood samples were collected from patients with carotid artery disease (CAD), upon patients’ informed consent, undergoing carotid endarterectomy surgeries at the Department of Vascular and Endovascular Surgery (Klinikum rechts der Isar, TUM). Blood samples were collected in EDTA tubes and kept at room temperature until further processing.

#### T cell labeling

Cells were stained using 1µM of either CellTracker™ Green CMFDA (C2925, ThermoFisher Scientific) or CellTracker™ Deep Red (C34565, ThermoFisher Scientific) diluted in serum-free culture medium for 45 minutes at 37°C. Afterward, cells were rinsed twice with PBS (10010023, ThermoFisher Scientific) and then stained with SYTO® 40 Blue Fluorescent Nucleic Acid Stain (S11351, ThermoFisher Scientific) diluted in serum-free culture medium for 45 minutes at 37°C.

#### T cell treatment with Pertussis Toxin and AMD-3100

After being labeled, T cells were treated with either 20nM Pertussis Toxin (PHZ1174, Gibco Life Technologies) or 1µM AMD-3100 (S8030, Selleckchem) for 1 hour at 37°C. Both Pertussis Toxin and AMD-3100 were diluted in serum-free medium to reach the final concentration.

### 2.5. Flow cytometry analysis

Plaques were digested with an enzymatic cocktail containing Advanced RPMI 1640 (12633-012, Gibco Life Technologies) supplemented with collagenase XI from Clostridium histolyticum at 1.25 mg/ml (Sigma-Aldrich, Saint-Loius, MS, USA, cat. C9697) and desoxyribonuclease I at 0.2 mg/ml (Sigma-Aldrich, cat. D5319). Isolated cells were strained through a 30µm filter (04-0042-2316, Sysmex) and red blood cells were lysed for 5 minutes at room temperature using ACK Lysis buffer (A1049201, Gibco Life Technologies). To block the FcR-involved unwanted staining, the cell suspension was incubated with Human TruStain FcX (Biolegend, Cat. 422301) and afterward, incubated with Zombie NIR™ Fixable Viability Kit (Biolegend, Cat. 423105) to assess live vs. dead status of the cells. Finally, the cell suspension was stained with monoclonal antibodies against CD45 (PE, 304058, Biolegend), CD3 (BV711, 300464, Biolegend), CD69 (BV510, 310936, Biolegend), CD4 (BV786, 566807, BD Bioscences) and CD8α (BUV395, 563795, BD Bioscences). Flow cytometry was performed using a Fortessa X20 flow cytometer (Becton Dickinson Biosciences, San Jose, CA, USA). Data were acquired with Diva software and analysed with FlowJo (Tree Star, Ashland, OR, USA).

### 2.6. Optical tissue clearing

For optical tissue clearing, plaques were fixed in 2% PFA for 48 hours at 4°C. Following permeabilization with 2% TritonX (1.12298.0101, Merck Millipore) and blocking using 5% BSA (3737.4, Carl Roth), 0.2% Triton X-100 (1.12298.0101, Merck Millipore) and Human TruStain FcX (Biolegend, Cat. 422301), plaques were stained with primary antibodies directed at CD31 (M0823, Dako and NB100-2284, Novus Biologicals) and CXCL12 (MAB350, R&D Systems). Alexa Fluor 488 (406416, Biolegend) and Alexa Fluor 594 (405326, Biolegend) were used as secondary antibodies. Finally, plaques were incubated with the clearing solution RapiClear 1.55 (SUNJin Lab) at RT until the sample resulted see-through. Plaques were then mounted using RapiClear 1.55 (SUNJin Lab) on glass-slides (22037246 Superfrost plus, ThermoFisher) using single sided sticky iSpacer 0.2mm (SUNJin Lab) to prevent tissue flattening.

After the tissue clearing and imaging protocol, plaques were washed with PBS (10010023, ThermoFisher Scientific) and were embedded in paraffin. Sequential cross-sections (5 μm thick) were made throughout the embedded atherosclerotic plaques. The sections were stained with Hematoxylin & Eosin (H&E) staining performed by routine methods and with primary antibodies directed at CD31 (NB100-2284, Novus Biologicals) and ACKR1 (566424, clone 2C3, BD Biosciences). Alexa Fluor 488 (406416, Biolegend) was used as secondary antibody.

### 2.7. Microscopy

For visualization of optically cleared human plaques, confocal- or two-photon laser scanning microscopy was utilized.

### 2.8 Statistics

For scRNA-seq of human carotid tissues DEGs were determined using a statistical threshold corrected for multiple testing using the false discovery rate (FDR) adjustment (FDR-adjusted P < 0.05; fold change 2).

All results are expressed as mean ± SEM. Normality of data obtained was examined using the Shapiro-Wilk normality test. A 2-tailed Student’s t-test, One-Way ANOVA or paired t-test were used to compare individual groups. Non-Gaussian distributed data were analyzed using a Mann–Whitney U test using GraphPad Prism version 6.00 for Windows (GraphPad Software). Probability-valuesLJ<LJ0.05 were regarded as significant. A detailed description of the methods and related references can be found in the Online Data Supplement.

## 3. Results

### 3.1. CD8^+^ T cells represent an abundant inflammatory cell population in human atherosclerotic plaques

We examined the presence of T cell populations in the cohort of patients from the Munich Vascular Biobank, using scRNA-Seq of advanced human atherosclerotic plaques (Fig. 1A). Our analysis identified a total of 15 populations of cells (Fig. 1B) of which 8 leukocyte cell clusters could be defined based on their expression of the pan-inflammatory marker gene CD45 (Fig. 1C and D). Three distinct T cell populations were identified (Fig. 1E), based on the expression of several marker genes including CD2, CD3D, CD3E and NKG7 (Fig. 1F). Our analysis showed that Cytotoxic CD8^+^ T cells (Fig.1G) were the most abundant hematopoietic cell population, comprising 13.8% of all cells analysed (Fig. 1H) and 50% of the CD45+ cell cluster population (Fig. 1I). These findings confirm that CD8^+^ T cells represent an abundant hematopoietic cell population when performing scRNA-Seq analyses of human atherosclerotic plaques^4, 5^.

**Figure 1.**
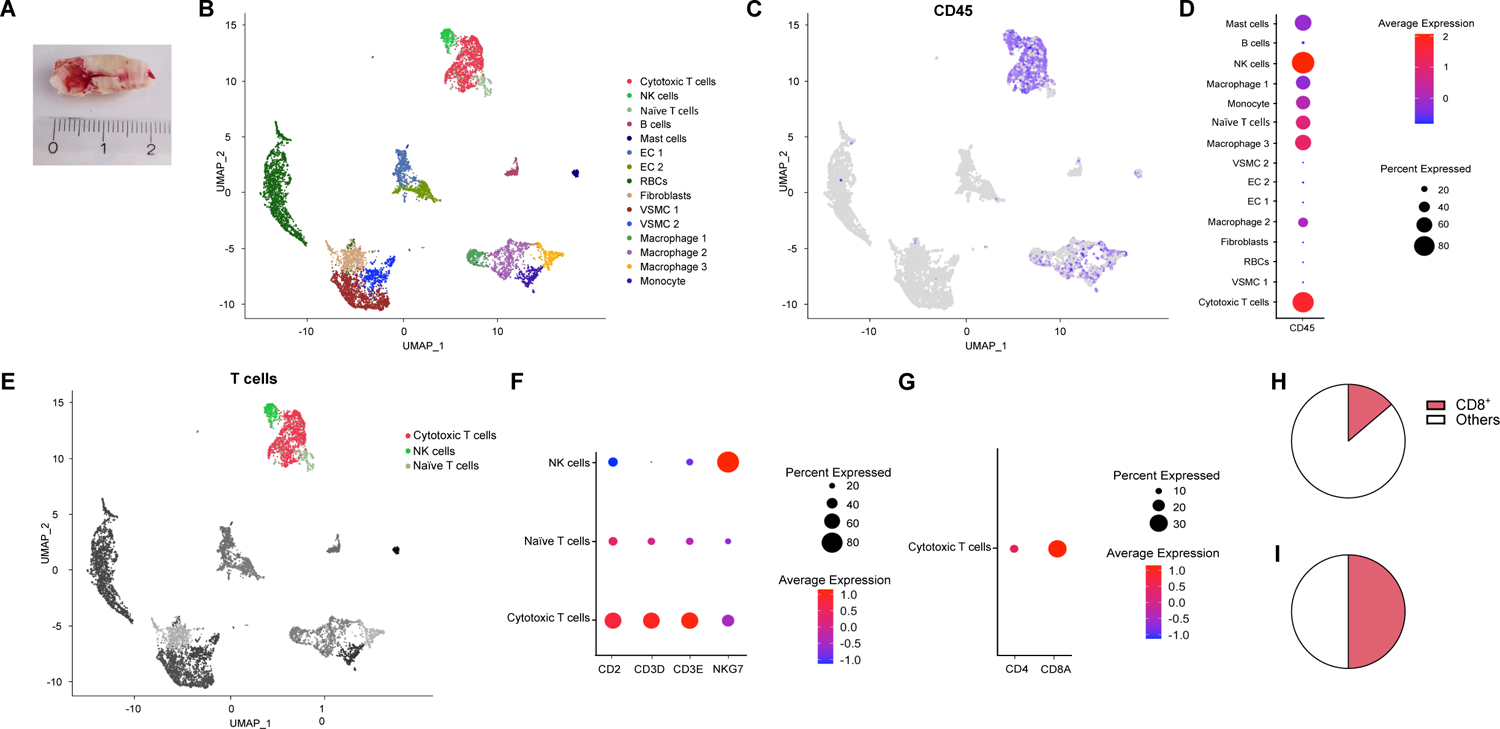
(A) Experimental setup: human plaque samples obtained from endarterectomy surgeries were digested, single viable cells were obtained, and single cell sequencing was performed. (B) UMAP visualization of clustering revealed 15 cell populations. Population identities were determined based on marker gene expression. (C) UMAP and (D) dot-plot visualization of CD45 clustering. (E) UMAP visualization of T cells clustering revealed 3 distinct populations. (F,G) Dot-plot of T cell clusters identifying genes. Percentage of CD8^+^ T cells and other T cells (H) among the total plaque population and (I) among the CD45^+^.

### 3.2. Quantitative evaluation of exogenous T cells infiltration in 3D *ex-vivo* co-cultured plaques

It was previously shown that plaque biopsies from patients affected by CAD, could be kept in culture for several days^18^. To evaluate whether patient-matching derived T cells could be co-cultured with human endarterectomies, we isolated CD8^+^ and CD4^+^ T cells from patient-matching blood samples (referred to as exogenous T cells) and stained them with a multi-factor staining strategy using different intracellular fluorescent cell tracker dyes combined with nuclear labels. The exogenous T cells were incubated in a petri-dish together with fresh patient-matching CEA biopsies that were processed in 1mm thick tissue slices and placed on a collagen sponge at the air-medium interface (Fig.2A). Two different plaque slices were kept in co-culture with exogenous T cells for 24 hours, then harvested and subsequently used for flow cytometry (FACS) and microscopic analysis respectively.

**Figure 2.**
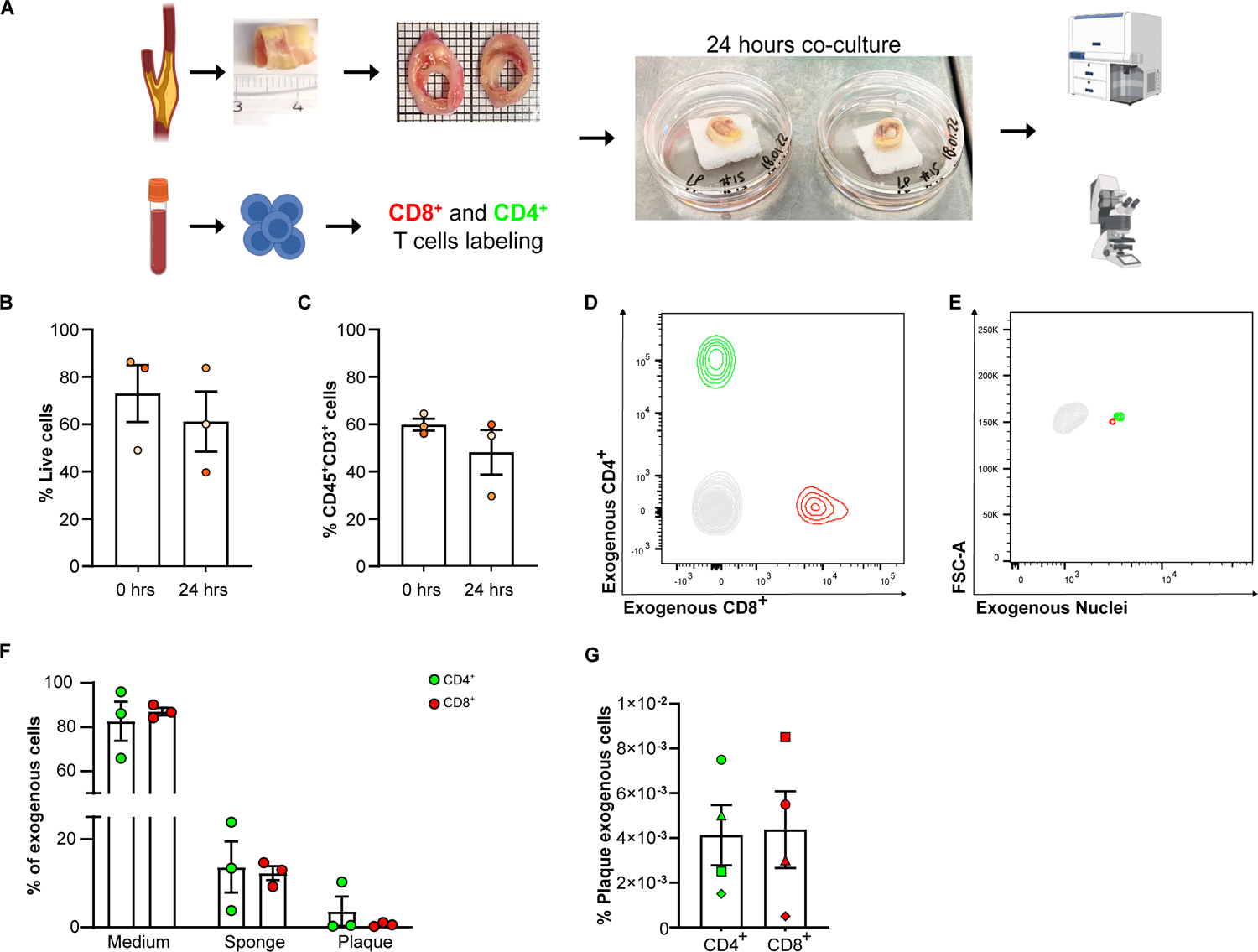
(A) Schematic representation of the 3D tissue-culture model of atherosclerosis. Atherosclerosis plaques from carotid endarterectomies were sliced in 1mm thick sections and T cells were isolated from patient-matching blood samples. T cells and atherosclerotic plaque slices were co-cultured for 24 hours and were processed for flow cytometry analysis and imaging respectively. Flow cytometry quantification of (B) total live cells and (C) total CD3+ cells in patient-matching fresh human atherosclerotic plaques (0hrs) as well as after co-culture with exogenous T cells infiltration in the plaque based on (D) Cell Tacker expression as well as (E) nuclear marker expression in CD45+CD3+ cells. (F) Distribution of exogenous CD4+ and CD8+ T cells in the different compartments of the culture (cell culture medium, sponge and plaque) calculated as fractions of the total amount of exogenous T cells in the culture. (G) Percentage of exogenous CD4+ and CD8+ T cells infiltrated in human atherosclerotic plaques after 24 hours of co-culture, calculated as fractions of the initial input of exogenous T cells (n=4, each shape represents a different patient). Values show mean ± SEM. Statistical significance was determined using Student’s t-test (two-tailed, unpaired).

Using flow cytometry, we found that plaques, when kept in co-culture with exogenous T cells for 24 hours, showed no difference in the total number of live cells when compared to control patient-matching plaques that were not kept in culture, but analysed on the day of the CEA (Fig. 2B, p=0.53). Interestingly, also the percentage of T cells in the plaque did not differ between plaques that were co-cultured with exogenous T cells and patient-matching plaques analysed on the day of the surgery (Fig. 2C, p=0.29). This demonstrates that neither the *ex-vivo* culture, nor the addition of exogenous T cells have a detrimental effect on the viability and composition of the plaques.

Next, we evaluated whether the exogenous T cells migrated into the plaque. Exogenous T cells were labelled using a multi-factor staining strategy: intracellular fluorescent staining (Cell Tracker^TM^) combined with a fluorescent nuclear staining (SYTO^®^) in order to distinguish them from the highly auto fluorescent plaque environment. Flow cytometry analysis (complete gating strategy in Supplemental Fig.1) showed that both exogenous CD8^+^ and CD4^+^ T cells, each labelled with a different fluorescent Cell Tracker dye could be found in the plaque after co-culture (Fig. 2D). The specificity of the two detected exogenous populations could be confirmed by the positivity for the nuclear labelling when compared to the endogenous cells in the plaque, which do not possess any nuclear staining (Fig.2E). Interestingly, we found that CD4^+^ and CD8^+^ T cells were equally distributed in the different culture compartments (Fig. 2F) and that both migrated into the plaques with comparable efficacy (Fig. 2G, p=0.88). These data indicate that migration of both T cell subtypes can be studied using the *ex-vivo* co-culture platform.

### 3.3. Mapping of exogenous T cells infiltration in *ex-vivo* co-cultured plaques

To visualize and map the location of the exogenous T cells infiltrated in the plaque, we applied optical tissue clearing using RapiClear 1.55, a water-soluble clearing reagent. Optical clearing enabled us to efficiently visualize the whole, multidimensional plaque specimen. The optical clearing procedure did not visibly affect the integrity of the tissues as only a marginal tissue deformation was observed between the plaques before and after the clearing protocol, which is in accordance with literature^19^ (Fig. 3A). Moreover, H&E staining of plaques after the optical tissue clearing protocol confirmed that the plaque maintained its structural and composition characteristics that were already present in the tissue before the clearing protocol and more broadly, can be typically found in human atherosclerotic lesions such as necrotic areas (Fig. 3B, 1) and areas of intraplaque angiogenesis (Fig.3B, 2).

**Figure 3.**
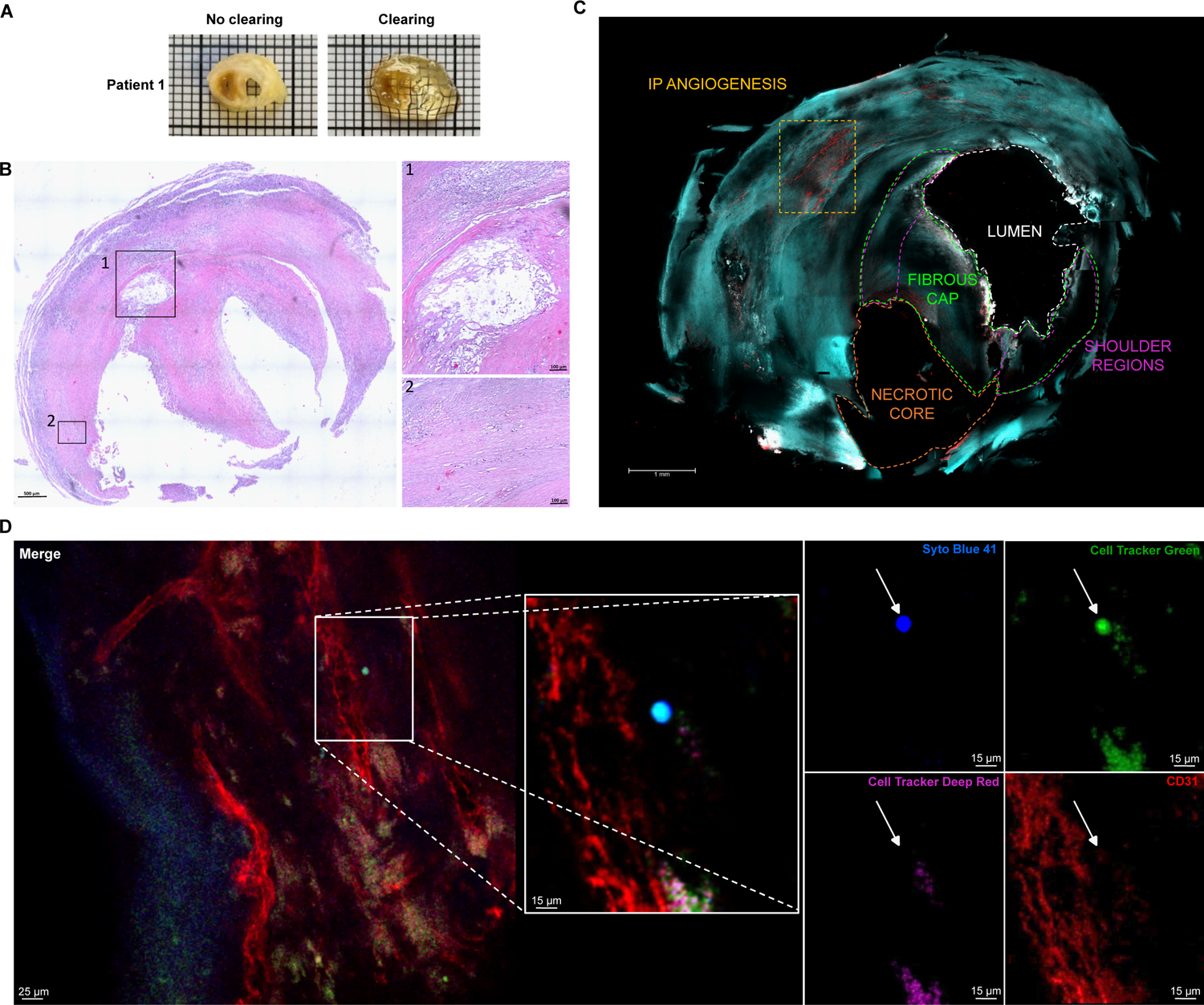
(A) Representative pictures of a human atherosclerotic plaque after 24 hours co-culture with patient-matching exogenous T cells without (left) and with (right) optical tissue clearing. (B) H&E staining of a human atherosclerotic plaque after the optical tissue clearing protocol. (C) Overview of an optically cleared human atherosclerotic plaque imaged using a confocal microscope, with the presence of Lumen, Fibrous Cap, Shoulder Regions, Necrotic Core and Areas of Intraplaque Angiogenesis. (D) Representative images of the multi-factor staining strategy used to discriminate between exogenous T cells and autofluorescence artifacts. Exogenous T cells were stained with Cell Tracker Green and Syto Blue 41 and are indicated by white arrows.

Using advanced optical microscopy modalities (confocal and/or two-photon laser scanning microscopy) combined with an automatic stage and automated stitching functions, we were able to visualize optically cleared plaques slices in full and create an overview in which the lumen, the necrotic core, shoulder region, fibrous cap and areas of intraplaque angiogenesis could be recognized (Fig. 3C).

Exogenous T cells were found in different parts of the plaque, for example in proximity of the shoulder regions (Supplemental Fig.2A) and in areas of intraplaque angiogenesis (Supplemental Fig.2B). The multi-factor staining strategy combined with optical clearing and microscopy allowed us to discriminate between exogenous T cells and auto fluorescence artefacts (Fig.3D). Round-shaped structures that were both positive for one of the exogenous intracellular markers and for the nuclear marker, with a size between 5 and 10 µm were regarded as infiltrated exogenous T cells (white arrows in Fig.3D). On the other hand, structures that showed a positive signal for both exogenous intracellular markers and the nuclear marker, and/or were not within the 5-10 µm size range were considered a consequence of tissue auto fluorescence and thus excluded from further analyses.

### 3.4. Chemokine receptors guide T cell infiltration in human atherosclerotic plaques

Chemokine receptors have been previously shown to play a role in T cells recruitment to the plaque^20, 21^. We evaluated the expression of chemokine receptors on T cell clusters present in human atherosclerotic plaques by analyzing the Munich Vascular Biobank scRNA-Seq dataset (Fig. 4A). We found that CXCR3, CXCR4, CXCR6 as well as CCR5 were highly expressed on T cells (Fig. 4A). Furthermore, this observation was particularly evident in cytotoxic CD8^+^ T cells. The expression pattern of these chemokine receptors was confirmed in various subpopulations of CD8^+^ and CD4^+^ T cells by analysing the human atherosclerotic plaques scRNA-Seq dataset from Slenders et al. (available on PlaqView platform^22^), which comprises several T cell sub-clusters^15^ (Supplemental Fig.3). Since CD8^+^ T cells are an abundant leukocyte population in human plaques^4, 5^, we next focused our investigation on CD8^+^ T cells exclusively.

**Figure 4.**
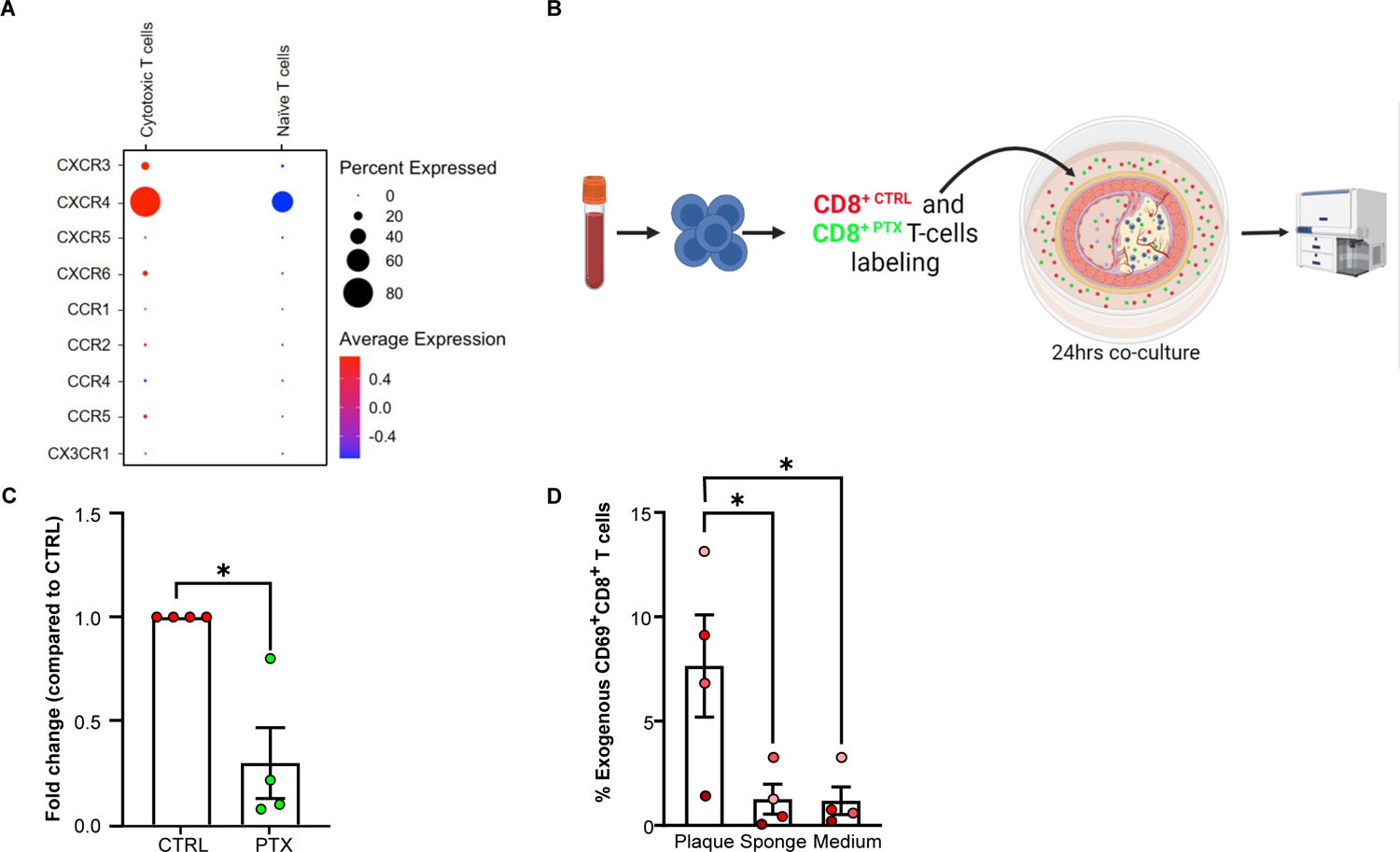
(A) Dot plots showing gene expression profiles of different chemokine receptors in T cells sub-clusters found in human atherosclerotic plaques. (B) Schematic representation of the experimental work-flow. (C) Flow cytometry analysis of plaque infiltrated CTRL and PTX-treated exogenous CD8^+^ T cells (n=4, each dot represents a different patient). (D) Flow cytometry analysis of the expression of CD69 on exogenous CD8^+^ T cells present in different compartments of the culture (n=4, each dot represents a different patient). Values show mean ± SEM. Statistical significance was determined using Student’s t-test (two-tailed, unpaired) and One-sample t-test. -p < 0.05.

To determine if exogenous CD8^+^ T cells infiltrate plaques through an active process, we used pertussis toxin (PTX), a commonly utilized inhibitor of chemotaxis via blockade of G protein-coupled receptors (GPCRs), that significantly impacts the motility of various immune cells, including T cells^23, 24^. Half of the isolated exogenous CD8^+^ T cells were treated with PTX, while the other half was treated with vehicle control (Fig. 4B). Following a 24-hour period of co-culture, we evaluated the exogenous T cell plaque content using flow cytometry and found that there was a dramatic reduction in the number of cells treated with PTX when compared to control cells (Fig. 4C, p=0.02). These data indicate that exogenous CD8^+^ T cells actively migrate into *ex-vivo* cultured atherosclerotic plaques and point towards a key role for chemokine receptors in this migratory process.

Subsequently, we evaluated if the exogenous T cells in the plaque can engage with other cell types through antigen-specific activation or if they are passive bystanders. To examine potential recent antigen encounter and activation, expression of CD69, a proxy marker for assessing T-cell responsiveness to antigen stimulus, was assessed. CD69 expression was measured through flow cytometry on the surface of exogenous CD8^+^ T cells that infiltrated into the cultured plaque and compared to its expression on exogenous CD8^+^ T cells present in the sponge and in the culture medium respectively. A significant increase in CD69^+^CD8^+^ exogenous T cells was observed in the plaque when compared to the sponge and the medium (Fig. 4D, p=0.01 and p=0.01 respectively). This data suggests that exogenous CD8^+^ T cells that infiltrate into cultured human plaques are activated in the tissue, thus confirming that exogenous CD8^+^ T cells are active players within the plaque environment.

### 3.5. CXCR4/CXCL12 axis is a master regulator of exogenous CD8^+^ T cell infiltration in human atherosclerotic plaques

Among the chemokine receptors expressed by T cells in the plaque, CXCR4 stood out as the most prominently expressed by CD8^+^ T cells (Fig. 4A). To test whether blocking CXCR4 on exogenous CD8^+^ T cells could impair their recruitment to the plaque, we used the FDA-approved CXCR4 antagonist AMD-3100 (Fig. 5A). Flow cytometry analysis revealed that treatment with AMD-3100 significantly reduced the migration of exogenous CD8^+^ T cells to the plaque (Fig. 5B). Interestingly, there were 3-fold less AMD-3100 treated exogenous CD8^+^ T cells in the plaque when compared to their vehicle-treated patient-matching control CD8^+^ T cells (Fig. 5B, p=0.01), suggesting that CXCR4 plays a major role in the recruitment of CD8^+^ T cells. To study the routes of T cell infiltration, we next investigated the presence and location of CXCL12, the unique ligand for CXCR4, in the plaque. Immunofluorescence staining of whole mount optically cleared plaques combined with two-photon laser scanning microscopy revealed that CXCL12 was present on endothelial cells (ECs), and in particular on ECs lining intraplaque neovessels (Fig. 5C and D). To validate this novel finding, we conducted an analysis of the two distinct populations of ECs, referred to as EC1 and EC2, present in the scRNA-Seq dataset from the Munich Vascular Biobank (Fig. 1B and Fig.5E).

**Figure 5.**
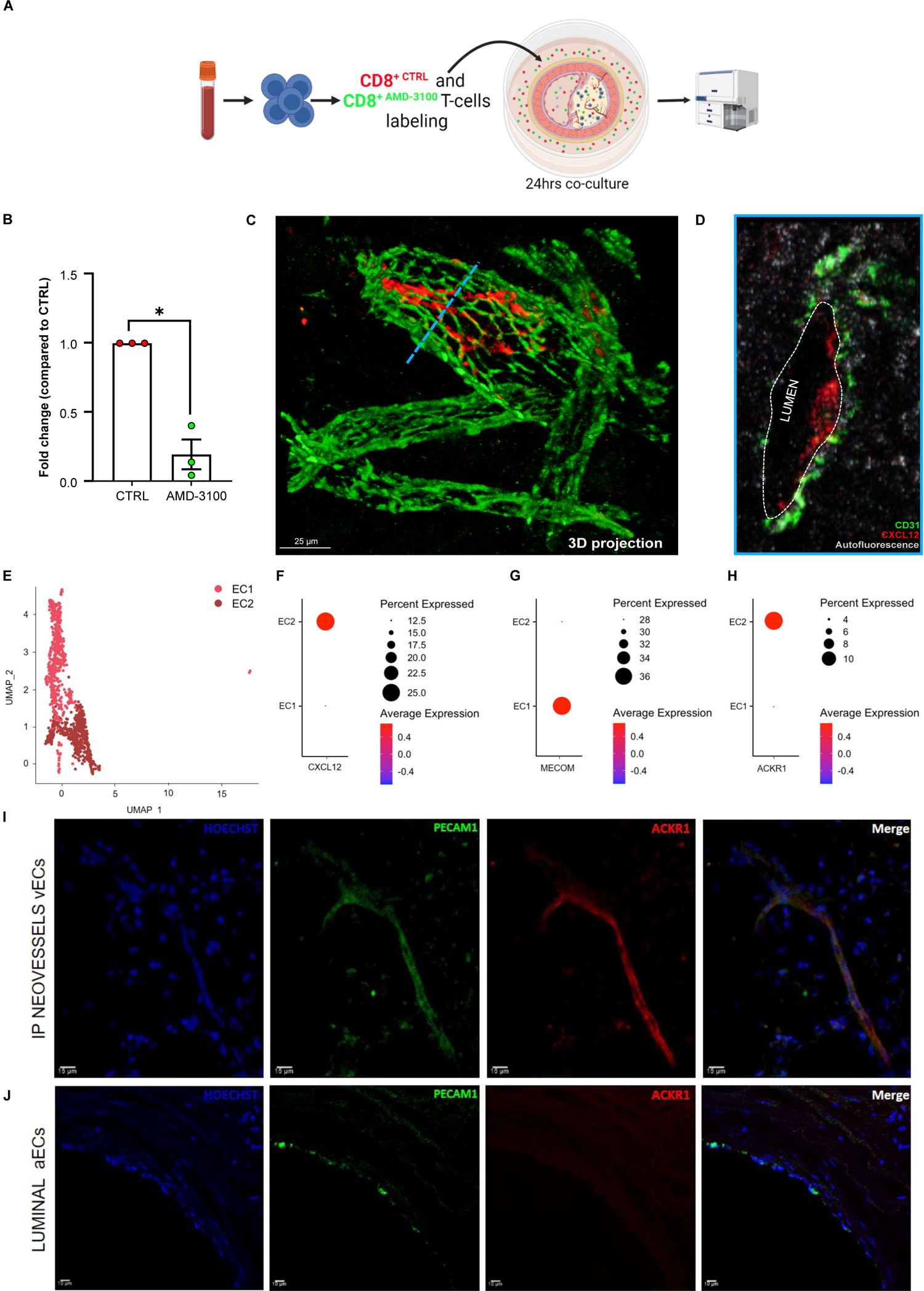
(A) Schematic representation of the experimental work-flow. (B) Flow cytometry analysis of plaque infiltrated CTRL and AMD3100-treated exogenous CDS+T cells (n=3, each dot represents a different patient). (C) 3D projection and (D) 2D image, corresponding to the transversal section outlined by the light-blue dotted line in panel C, of intraplaque neovessels expressing CXCL12 in human atherosclerotic plaque (CD31 in green, CXCL12 in red and tissue autofluorescence in grey). (E) UMAP visualization of endothelial cells clustering revealed 2 distinct populations, ECl and EC2. Dot-plot visualization of (F) CXCL12 and (G) arterial and (H) venous identifying genes. (I) Immunohistochemistry staining of intraplaque neovessels vECs and (J) lumen aECs in advanced human atherosclerotic plaques (Nuclei in blue, PECAMl in green and ACKRl in red). Values show mean ± SEM. Statistical significance was determined using One-sample t-test (two-tailed, unpaired). *p < 0.05.

Interestingly, our findings revealed that CXCL12 was highly expressed in the EC2 population (Fig. 5F). Further characterization of ECs showed that EC1 exhibited expression of several gene markers associated with arterial ECs (aECs) (MECOM, GJ4, GJ5, GATA 2, Fig. 5G and Supplemental Fig.4A), while EC2 displayed gene markers typically found in venular ECs (vECs) (ACKR1, NR2F2, PLVAP, Fig.5H and Supplemental Fig. 4A). These findings were supported by the analysis of the publicly available dataset of human atherosclerotic plaques^15^ described above (Supplemental Fig.4B and C). Therefore, we classified EC1 as luminal aECs and EC2 as intraplaque neovessels vECs. To further validate these findings, immunohistochemistry staining of advanced human atherosclerotic plaques was performed, confirming that the ECs lining the intraplaque neovessels exhibited positive staining for the pan-endothelial cell marker PECAM1, as well as the vECs marker ACKR1^25^ (Fig. 5I). Conversely, ECs forming the lumen of the carotid vessel showed no expression of ACKR1 (Fig. 5J). Altogether, these data demonstrate that ACKR1^+^ vECs of the intraplaque neovessels highly express CXCL12, providing an explanation for its abundant presence in these vessels.

### 3.6 Intraplaque neovessels represent a route for T cells entry into the plaque

The expression of CXCR4 on T cells and the presence of CXCL12 on vECs lining intraplaque neovessels indicate that these vessels may provide a potential pathway for the recruitment of T cells in human atherosclerosis. Since intraplaque angiogenesis typically occur in advanced atherosclerotic plaques, we further examined whether exogenous CD8^+^ T cells infiltrate into the plaque through this route. Our analysis confirmed the presence of exogenous CD8^+^ T cells inside the intraplaque neovessels, as well as in their proximity (Fig. 6A and B), indicating that infiltration through intraplaque neovessels is a possible route for T cell entry into the plaque. Overall, these findings suggest that exogenous CD8^+^ T cells can be recruited via intraplaque neovessels and engage with endothelial cells on the luminal side of intraplaque neovessels via CXCR4/CXCL12 interaction.

**Figure 6.**
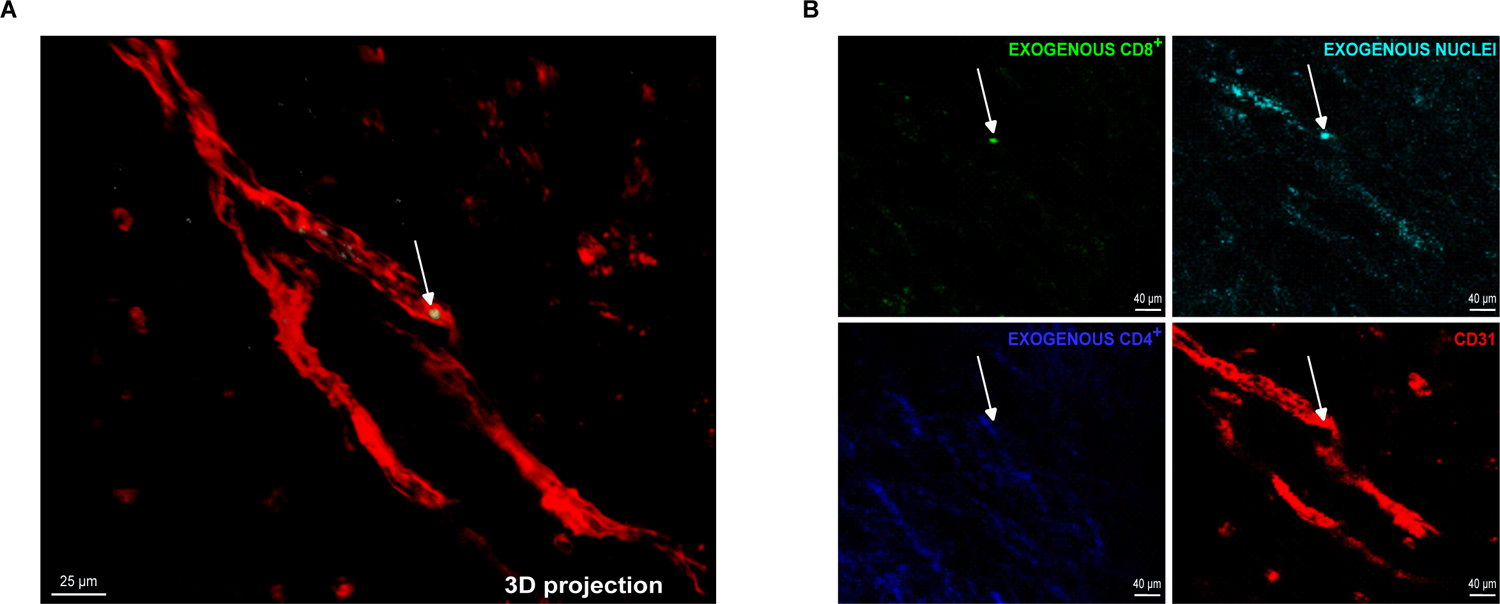
(A) 3D projection and (B) 2D images of exogenous CD8^+^ T cells within intraplaque neovessels (CD31 in red, exogenous nuclei in cyan, exogenous CD4^+^ in blue and exogenous CD8^+^ in green).

## 4. Discussion

In this study, we established a 3D tissue-culture model to study leukocyte recruitment to human atherosclerotic plaques. We show that human plaques derived from patients affected by CAD can be successfully co-cultured with patient-matching T cells and this co-culture model can be utilized to study leukocyte recruitment and accumulation in the plaque. Moreover, we demonstrated that this is not a passive accumulation but rather an active recruitment. We showed that CD8^+^ T cells utilize chemokine receptors to infiltrate atherosclerotic plaques, and in particular, we identified CXCR4 as the main chemokine receptor involved in this process. Utilization of optical tissue clearing combined with advanced microscopy modalities enabled us to localize CXCL12, the ligand of CXCR4, on ECs lining intraplaque neovessels, as well as exogenous CD8^+^ T cells within or in their vicinity, identifying a new possible explanation for the detrimental role played by CXCL12 in atherosclerosis.

Animal models are commonly used for atherosclerosis research^26–28^. However, the most widely used mouse models for atherosclerosis (such as ApoE^-/-^ or Ldlr^-/-^) only emulate the early stages of the disease, thereby impeding the ability to replicate the full complexity of the human pathology^26^. In fact, unlike human atherosclerosis, mouse lesions do not progress toward an unstable phenotype. For example, they do not physiologically develop intraplaque angiogenesis and hemorrhage, typical features of advanced human plaques^2, 29–32^. These limitations are one of the contributing factors to why many of the preliminary therapeutic candidates fail at a later stage of drug development. For this reason, there is a strong need to develop new scientific tools that more accurately mimic the human pathological condition.

Although in the past years, several 2D and 3D *in vitro* models of atherosclerosis have been developed^33–35^, these models are most often composed by only one or two cellular types and as such lack a relevant patho-physiological structure. More recently, Zhang *et al.* created a 3-layer Tissue-Engineered Blood Vessel (TEBV) composed of human ECs, SMCs and fibroblasts^36^. The authors observed changes in endothelial activation and permeability as well as in monocyte accumulation and foam cell formation in the vessel wall upon pro-inflammatory treatment^36^. While this study recapitulates the early atherosclerotic disease stage, it lacks a disease progression toward plaque destabilization. To overcome these issues, it was previously shown that biopsies from CAD patients could be kept alive for several days when cultured *ex-vivo*^18^. Here, we show that freshly isolated human plaques remain viable in *ex-vivo* culture and can be co-cultured with patient-matching T cells to study their infiltration and exact location in the plaque.

Following the traditional belief that macrophages derived from infiltrating monocytes and foam-cells dominate the atherosclerotic plaque landscape, previously developed *in vitro* models of atherosclerosis focused on the recruitment of monocytes to the plaque and their interaction with endothelial cells^37, 38^. However, as it has been shown that T cells represent a prominent cellular component in human plaques ^4, 5, 15^, we have developed a model to study T cell recruitment and localization in human atherosclerosis. Importantly, by utilizing patient-matching T cells, we were able to circumvent inter-patient variability and prevent the introduction of undesired immune responses. This model enables to quantitatively evaluate the recruitment of different T cells subtypes to the plaque via flow cytometry, as well as qualitatively assess structural properties such as the 3D location of recruited T cells. The combination of optical tissue clearing, a multi-factor staining strategy, and state-of-the-art microscopic modalities such as two-photon- and confocal laser scanning microscopy, enabled us to overcome the hurdle of imaging within the highly autofluorescent and strongly light scattering environment of human atherosclerotic plaques. Combined with the co-culture of plaques and patient-matching T cells, microscopy of optically cleared plaque allowed us to spatially localize exogenous T cells in the plaque and evaluate their infiltration in the tissue with µm precision in three dimensions. As such this model holds potential to be utilized as a drug testing platform for evaluation of different treatments on the recruitment of T cells as well as their effects on the whole plaque environment.

Chemokine receptors and their ligands play a critical role in the recruitment of T cells^21, 39^ and other inflammatory cells^40–42^ to atherosclerotic plaques. Recent scRNA-Seq studies showed that in human plaques, T cells overexpress several chemokine receptors^4, 15^. In our model, blocking these receptors by using PTX, significantly reduces CD8^+^ T cell infiltration into the plaque. Specifically, blocking CXCR4, one of the most highly expressed chemokine receptors, results in a drastic reduction of CD8^+^ T cell infiltration. Interestingly, large genome-wide association studies have linked the expression of CXCL12, the sole ligand for CXCR4, to CVDs^43–47^. Studies investigating the source of CXCL12 in atherosclerosis have been conducted in both human and mouse. One study using an immunohistochemistry approach failed to identify the source of CXCL12 in human atherosclerotic lesions^48^, likely due to technical issues related to the critical steps of fixation, permeabilization and washing^49^. Another study showed that, in mice, arterial endothelial cells produce CXCL12, which accelerates atherosclerosis formation^50^. However, due to technical issues in human samples and the absence of advanced plaques in mouse models, the main source of CXCL12 in atherosclerosis may have been missed in these studies. Utilizing advanced imaging techniques in combination with an optical tissue clearing strategy to examine whole human atherosclerotic plaques enabled us to identify the presence of CXCL12 on the luminal surface of intraplaque neovessels, revealing a new insight into its mechanism of action in atherosclerosis. Immunohistochemical staining and scRNAseq analysis provided additional evidence indicating that vECs from intraplaque neovessels are the primary source of CXCL12 in atherosclerotic lesions. Specifically, our findings suggest that CXCL12 mainly plays its role in intraplaque neovessels by providing a route for T cell extravasation and plaque infiltration. This conclusion is supported by the co-localization of CXCL12 and exogenous CD8^+^ T cells in close proximity and within intraplaque neovessels.

The ex-vivo 3D human atherosclerosis tissue-culture model offers several benefits for studying T cell recruitment; however, a potential drawback is the absence of flow, which means that exogenous T cells may not completely follow their natural recruitment pathway via the luminal side. Despite this limitation, our study clearly demonstrated that T cells can also be recruited by intraplaque neovessels. Furthermore, our findings indicate that exogenous T cells are actively recruited and become activated once inside the plaque, further highlighting the significance and complexity of this newly developed model. Moreover, while the sample sizes are limited and the methods are challenging, this method can provide details on specific interactions and complement genomic studies.

In conclusion, the here presented 3D tissue-culture model of atherosclerosis bridges the gaps that are currently present between pre-clinical studies and their translation into clinical practice and holds potential as a drug screening platform.

## Acknowledgements

Illustrative figures were created with BioRender.com.

## Supplemental Material

Supplemental Methods Figure S1, S2, S3 and S4

Table S1

References 1 and 2

