## Supplementary material for "Investigating T cell Recruitment in Atherosclerosis using a novel Human 3D Tissue-Culture Model reveals the role of CXCL12 in intraplaque neovessels": Online data supplement

A) Detailed Methods
B) Supplemental Figures and Tables
C) References

### A) Detailed Methods

***Single cell RNA sequencing (scRNA-seq)***

*Tissue Dissociation*: Plaques were minced and processed using the Multi Tissue Dissociation Kit 2 from Miltenyi Biotech (catalog number 130-110-203), along with the GentleMACS Dissociator (catalog number 130-093-235) and GentleMACS C tubes (catalog number 130-096-334), all following the instructions provided by the manufacturer. The resulting cell suspension was strained through a 70-µm filter and underwent Dead Cell Removal (Miltenyi Biotech, catalog number 130-090-101) using MS Columns (catalog number 130-640-042-201). Finally, the cells were suspended in PBS containing 0.04% BSA.

*Single-cell capture and library preparation*: Cells were loaded into a 10x Genomics microfluidics Chip G and encapsulated with barcoded oligo-dT-containing gel beads using the 10x Genomics Chromium Controller. Gel Beads-in-emulsion (GEM) cleanup, cDNA Amplification and 3´Gene Expression Library Construction was performed according to the manufacturer´s instructions (CG000204 Rev D). The resulting libraries from individual samples were multiplexed into a single lane and sequenced using an Illumina NovaSeq6000 instrument.

*ScRNA-seq analysis*: R package Seurat (version 4.1.1) was used in R Studio (version 1.4.1717) for scRNA-seq analysis. 18 samples of 9 patients are included in the final analysis (9 early stage samples and 9 advanced stage samples). Genes were excluded for downstream analysis if expressed in fewer than 5 cells and, in each Seurat object, cells with maximum mitochondrial reads >15%, maximum UMIs more than 20,000, minimum genes less than 100 and maximum genes more than 3000, were filtered out. To obtain the scores of cell cycle phases, including S phase and G2M phase, CellCycleScoring function in Seurat was applied and SCTransform normalization workflow was adopted to mitigate possible variations. UMAP (Uniform Manifold Approximation and Projection) combined with FindAllMarkers package was used to convert cells into a two-dimensional map and detect the main features of each cluster with default parameters.

**T cells isolation, labelling and treatment**

*Blood processing for simultaneous CD4+ and CD8+ isolation*: Human peripheral blood mononuclear cells (PBMCs) were isolated from the blood samples using a density gradient approach with Histopaque-1077 (SigmaAldrich) following manufacturer's instructions. CD4+ and CD8+ were then isolated starting from the PBMCs using CD4+ T Cell Isolation Kit (130-096-533, Miltenyi Biotec) and CD8+ T Cell Isolation Kit (130-096-495, Miltenyi Biotec) following manufacturer's specification. For the isolation LS columns (130-042-401, Miltenyi Biotec) and MidiMACS™ Separator were used (130-042-301, Miltenyi Biotec). Cells were then counted and equal numbers of CD4+ and CD8+ T cells were incubated with each plaque slice.

*Blood processing for CD8+ isolation*: For the isolation of solely CD8+ T cells, whole blood was processed using EasySep™ Direct Human CD8+ T Cell Isolation Kit (19663, StemCell Technologies) and "The Big Easy" EasySep™ Magnet (18001, StemCell Technologies) following manufacturer's instructions. Cells were then counted and incubated with each plaque slice.

**Microscopy**

For confocal microscopy, a Leica TCS SP8 3X (Leica Microsystems, Germany) equipped with a UV laser, a freely tunable white light laser, hybrid diode detectors and a 16X0.6 IMM-objective (designed for optical clearing; WD 2.5mm) were applied. The spectral excitation and emission parameters were optimized for sequential acquisition of each dye in order to maximize signal collection while avoiding in-between-channel spillover: Syto-40 Blue: excitation 405nm, detection 420-465nm; Cell Tracker Green 488: excitation 496nm, detection 500-540nm; Cell

Tracker Deep red: excitation 635nm, detection 655-700nm; Alexa Fluor 594: excitation 590nm, detection 610-650nm. Autofluorescence signals of plaque components were detected in a dedicated channel (excitation; 560nm, emission; 575-620nm). In combination with the background autofluorescent signal present in both the Syto-40 Blue and Celltracker green spectral channels, a mixed color autofluorescence is generated in overlay images thereby enhancing contrast between autofluorescent and exogenous stainings. For more insight in plaque structure in part of the samples, two-photon laser scanning microscopy was utilized to visualize the endogenous fluorescence of cleared human plaques. The latter was performed with a Leica SP5II MP system coupled to a pulsed Ti:Sa Laser (Spectra Physics MaiTai DeepSee, Newport Mannheim, Germany), tuned at 840nm, a 20X1.00 water dipping (WD 2mm) objective, 4 hybrid diodes. Imaging was performed non-sequentially and autofluorescence signals from plaque and detection of exogenous markers were detected in all four channels (410-425nm; 430-500nm; 550-570nm; 675-660nm). Second harmonics Generation (SHG) from (mainly) collagen was collected in the first spectral channel at 420nm. Z-stacks of 250-800μm were collected with a 2-4μm step-size. Raw pictures were processed and deconvolved using Leica Application Suite X with 3D-advanced and lightning plugins (version 3.5.1, Leica Microsystems, Germany).

For ACKR1 staining, the antibody 566424, clone 2C3, BD Biosciences was used[1].

**B) Supplemental Figures and Tables**

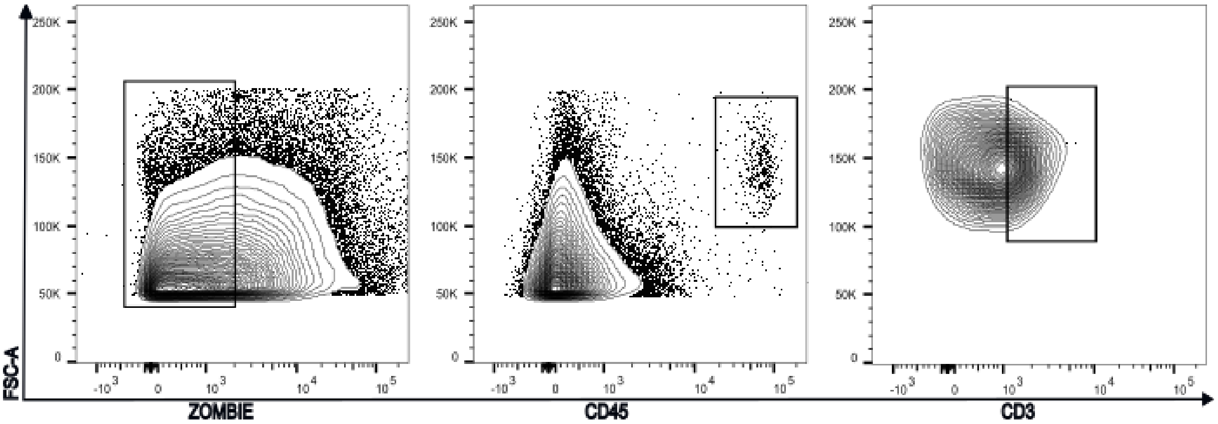


**Supplemental Figure 1.** Representative flow cytometry gating strategy for the analysis of exogenous T cells infiltration in the plaque based on Zombie live/dead staining, CD45 expression as well as CD3 expression.

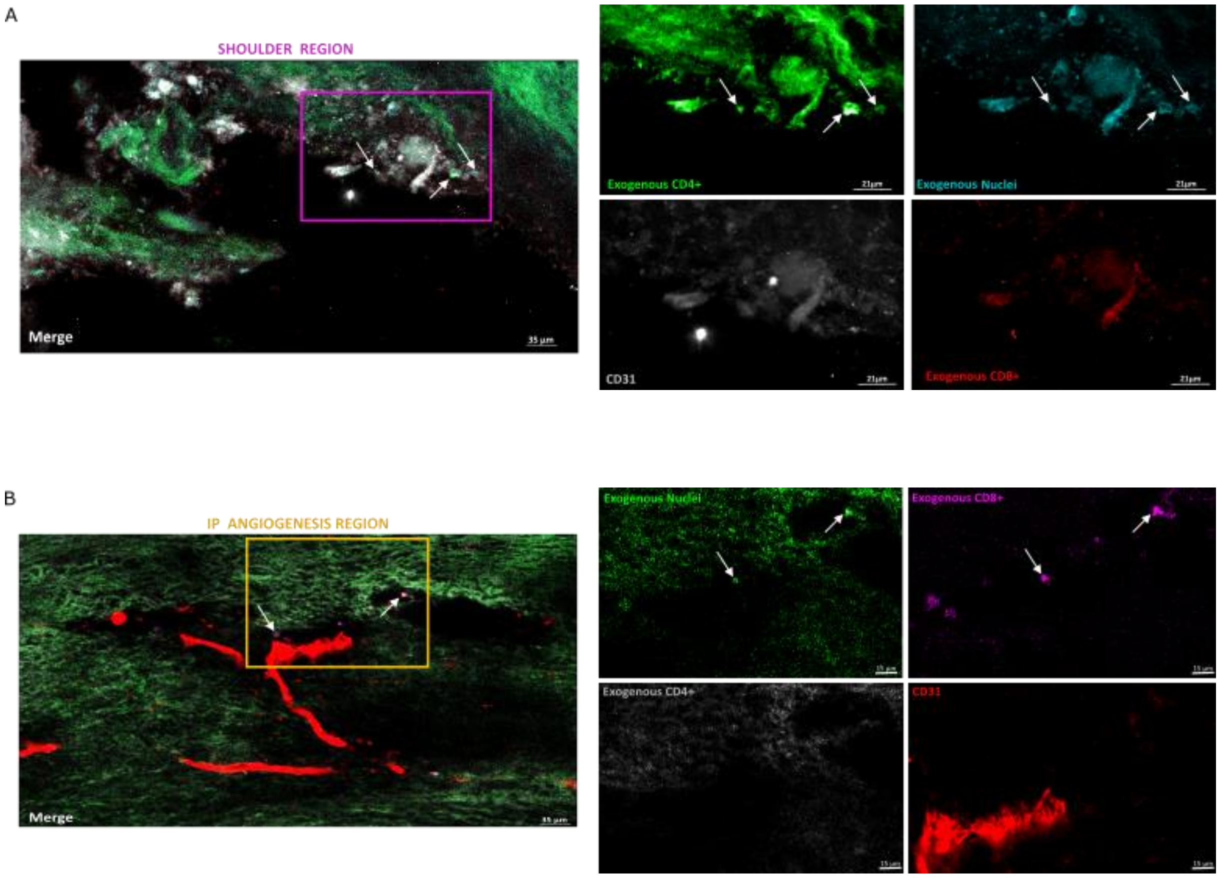


**Supplemental Figure 2.** Analysis of (A) a shoulder region area and (B) an area of intraplaque angiogenesis in which several exogenous $CD4^+$ T cells and exogenous $CD8^+$ T cells could be found, as confirmed by double staining of Cell Tracker and nuclear marker. Exogenous $CD4^+$ and $CD8^+$ T cells are indicated with white arrows.

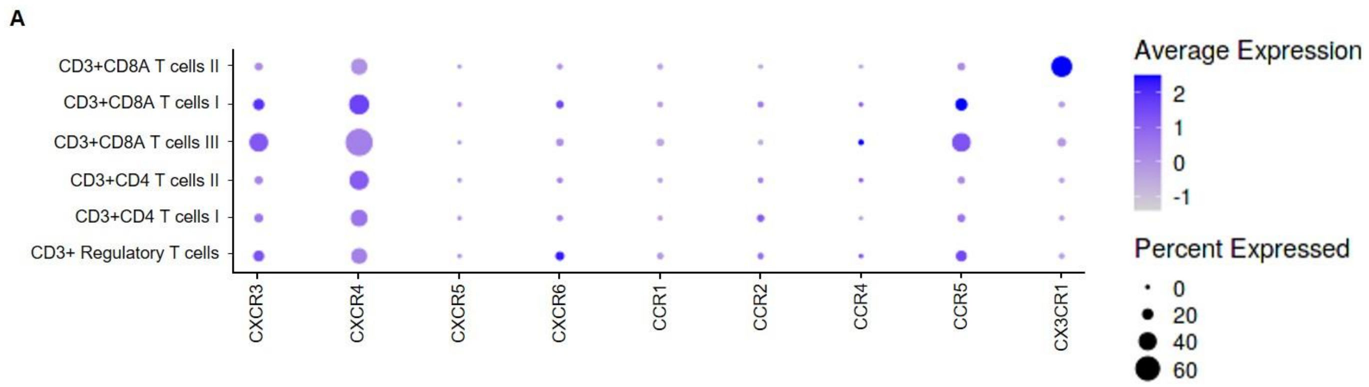


**Supplemental Figure 3.** (A) Dot plots showing gene expression profiles of different chemokine receptors in T cells sub-clusters found in human atherosclerotic plaques[2] (http://plaqviewv2.uvadcos.io/) .

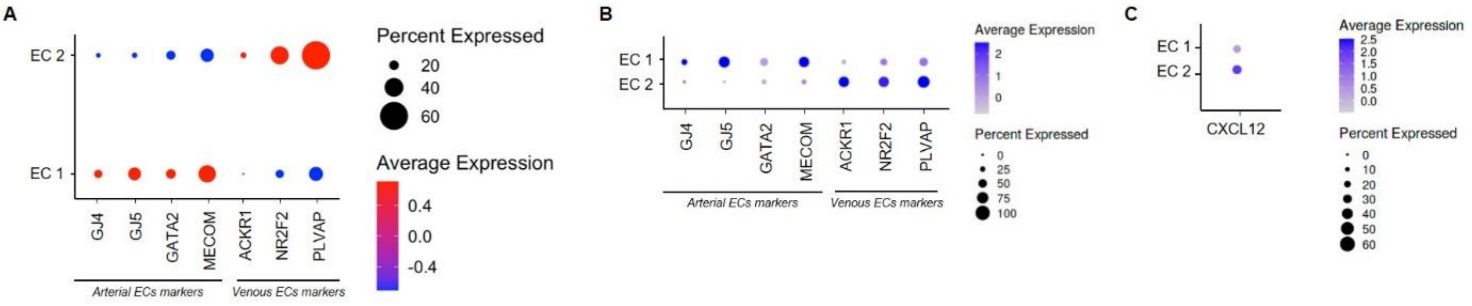


**Supplemental Figure 4.** Dot-plot visualization of several arterial and venous endothelial cells identifying genes from (A) the the Munich Vascular Biobank and (B) Slenders et al.[2] using the http://plaqviewv2.uvadcos.io/. (C) Dot-plot visualization of CXCL12 gene in the dataset from Slenders et al.[2] using the http://plaqviewv2.uvadcos.io/ online platform.

| Age | Gender | Symptoms | Degree of stenosis (NASCET) | Hypertension | Diabetes | Nicotine | Dyslipidemia | CAD | MI | AF | PAD | COPD | Dialysis | Height (cm) | Weight (Kg) | BMI | PFI | Statins | Insulin |
|---|---|---|---|---|---|---|---|---|---|---|---|---|---|---|---|---|---|---|---|
| 84 | m | a | 70 | yes | no | ex | no | no | no | no | no | no | no | 163 | 58 | 21,8 | yes | yes | no |
| 75 | m | s | 70 | yes | yes | no | no | yes | yes | no | no | no | no | 170 | 80 | 27,7 | yes | no | no |
| 79 | f | s | not known | yes | no | no | no | no | no | no | no | no | no | 162 | 75 | 28,6 | yes | alternativ | no |
| 74 | m | a | 90 | yes | no | no | yes | yes | yes | no | no | no | no | 170 | 90 | 31,1 | yes | yes | no |
| 76 | m | a | 80-90 | yes | no | ex | yes | no | no | no | no | no | no | 165 | 80 | 29,4 | yes | yes | no |
| 74 | m | a | 80-90 | yes | no | no | yes | no | no | no | no | no | no | 172 | 95 | 32,1 | yes | yes | no |
| 75 | m | s | 80 | yes | no | no | yes | no | no | no | no | no | no | 172 | 62 | 21,0 | yes | yes | no |
| 73 | m | a | not known | yes | no | ex | yes | yes | no | yes | no | no | no | 169 | 65 | 22,8 | yes | yes | no |
| 85 | f | a | 70 | yes | yes | no | yes | yes | yes | no | no | no | no | 143 | 47 | 23,0 | yes | yes | no |

**Supplemental Table 1.** Patient´s information.
